# Alpha-synuclein-induced nigrostriatal degeneration and pramipexole treatment disrupt frontostriatal plasticity

**DOI:** 10.1101/2024.02.17.580817

**Authors:** Sarah Chevalier, Mélina Decourt, Maureen Francheteau, Anaïs Balbous, Pierre-Olivier Fernagut, Marianne Benoit-Marand

## Abstract

**BACKGROUND:** Parkinson’s disease is characterized by the degeneration of *substantia nigra pars compacta* (SN*c*) dopaminergic neurons, leading to motor and cognitive symptoms. Numerous cellular and molecular adaptations due to the degenerative process or dopamine replacement therapy (DRT) have been described in motor networks but little is known regarding associative basal ganglia loops.

**OBJECTIVE:** To investigate the contributions of nigrostriatal degeneration and pramipexole (PPX) on neuronal activity in the orbitofrontal cortex (OFC), frontostriatal plasticity and markers of synaptic plasticity.

**METHODS:** Bilateral nigrostriatal degeneration was induced by viral-mediated overexpression of human mutated alpha-synuclein in the SNc. Juxtacellular recordings were performed in anesthetized rats to evaluate neuronal activity in the OFC. Recordings in the dorsomedial striatum (DMS) were performed and spike probability in response to OFC stimulation was measured before and after a high frequency stimulation (HFS). Post-mortem analysis included stereological assessment of nigral neurodegeneration, BDNF and TrkB levels.

**RESULTS:** Nigrostriatal neurodegeneration led to altered firing patterns of OFC neurons that were restored by PPX. HFS of the OFC led to an increased spike probability in the DMS, while dopaminergic loss had an opposite effect. PPX led to a decreased spike probability following HFS in control rats and failed to counteract the effect of dopaminergic neurodegeneration. These alterations were associated with decreased levels of BDNF and TrkB.

**CONCLUSIONS:** Both nigral dopaminergic loss and PPX concur to alter fronstostriatal transmission, precluding adequate information processing in associative basal ganglia loops as a gateway for the development of non-motor symptoms or non-motor side-effects of DRT.

## INTRODUCTION

Parkinson’s disease (PD) is characterized by the degeneration of several neuronal ensembles including dopaminergic neurons in the *substantia nigra pars compacta* (SN*c*), and by the occurrence of cytoplasmic inclusions containing misfolded alpha-synuclein among other aggregated components. In addition to the cardinal motor symptoms of the disease, several non-motor symptoms that are often underdiagnosed are frequently observed at disease onset. These include apathy, anhedonia and subtle executive dysfunction, as evidenced by attentional deficits, memory complaints and decreased cognitive flexibility [1–3]. Some of these cognitive impairments have been associated with reduced [^18^F] fluorodopa uptake in the caudate nucleus and frontal cortex [4], reflecting the ongoing loss of dopaminergic innervation of the medial prefrontal and orbitofrontal cortices arising from both the ventral tegmental area (VTA) and to a lesser extent form the SNc [5, 6].

The consequences of dopaminergic denervation on the functioning and information integration in basal ganglia motor circuits have been largely investigated in animal models of PD and have revealed that loss of dopaminergic innervation leads to altered activity in both the motor cortex and dorsolateral striatum. Loss of midbrain dopaminergic neurons leads to decreased excitability [7] as well as reduced firing rate of pyramidal neurons in the primary motor cortex in rats [8] and non-human primates [9], although such effects seem to be influenced by the type of dopamine depletion (acute vs chronic) and species used to model PD [10]. At the striatal level, dopamine depletion leads to synaptic remodeling, together with altered morphology and activity of striatal output neurons [11]. Furthermore, cortico-striatal transmission is also markedly altered following nigrostriatal dopamine depletion, as evidenced by impairments of Hebbian plasticity (long-term potentiation and long-term depression) (for review, see [12]). Whether these cortical and striatal functional alterations demonstrated in motor circuits similarly operate in associative domains of the basal ganglia remain currently poorly known.

The introduction of dopamine replacement therapy alleviates most of the motor deficits and normalizes to some extent the abnormal activity of cortical and subcortical nuclei but its long-term use leads to the development of maladaptive forms of synaptic plasticity that underlie the expression of motor-side effects such as dyskinesia or non-motor side-effects such as impulse control disorders [13–15]. Based on the functional organisation of the basal ganglia, both motor and non-motor side effects of dopaminergic treatments have been theorized to represent a pathological continuum of abnormal dopaminergic stimulation, with possible shared underlying molecular and cellular mechanisms [15].

In this study we aimed at determining if circuit dysfunctions described in motor cortico-striatal loops following nigrostriatal degeneration and dopamine replacement therapy find their counterparts in associative fronto-striatal loops. To this end, we used a rat model of PD based on viral-mediated expression of alpha-synuclein in the SNc to investigate the consequences of the degeneration of nigral dopaminergic neurons and of chronic treatment with the D3/D2 agonist pramipexole (PPX) on neuronal activity and plasticity in fronto-striatal circuits. To this end, we recorded the spontaneous activity of orbitofrontal cortex (OFC) neurons and evaluated fronto-striatal plasticity, as assessed by recording the response of neurons in the dorsomedial striatum to OFC stimulations, before and after a high frequency stimulation protocol known to induce a long-term potentiation [16].

## METHODS

### Animals

All experiments were approved by the local ethical committee (Comethea Poitou-Charentes C2EA-84, study approval #24114-2020021310245838) and performed under the European Directive (2010/63/EU) on the protection of animals used for scientific purposes. Male Sprague Dawley rats (N=76, 175g, Janvier, France) were housed on a reversed 12h cycle. Food and water were available *ad libitum*. Rats were segregated into four groups: sham treated with saline (GFP SALINE, N=17), sham treated with PPX (GFP PPX, N=20), lesioned treated with saline (alpha-SYN SALINE, N=19) and lesioned treated with PPX (alpha-SYN PPX, N=20).

### AAV-alpha-synuclein-mediated lesion

Rats were anesthetized with isoflurane (2% at 1.5 L/minute), vitamin A was applied on the rat’s eyes to prevent corneal dryness and 1 mL NaCL 0.9% was injected subcutaneously to prevent dehydration. Pre-surgical analgesia (ketoprofen, 10 mg/kg, i.p.) and local analgesia (xylocaine gel 2%) were performed following vetidine disinfection. Rats were placed in a stereotaxic frame (Kopf Instruments®, Tujunga, CA, USA) and after skin incision, the skull was drilled to perform two bilateral injections in the SNc (in mm from bregma and dura, AP: -5.1 and -5.6; ML: +/-2.2 and +/-2 ; DV: -8) with 1µL of AAV2 expressing GFP (green fluorescent protein, CMVie/SynP-GFPdegron-WPRE, 3.7 10^13^ gcp/mL) or AAV2 expressing human A53T alpha-synuclein (CMVie/SynP-synA53T-WPRE, 5.2 10^13^ gcp/mL) at 0.2µL/min as previously described [17–19]. Post-surgical analgesia was performed with ketoprofen (10 mg/kg/day, i.p.) the day after surgery and during three days if necessary.

### Treatment

Eight weeks after stereotaxic surgery, rats received either saline (NaCl 0.9%) or PPX 0.3mg/kg/day (Sequoia Research Products, Berkshire, UK) prepared daily in sterile saline solution. Both were administrated subcutaneously for 10 days [19].

### Electrophysiological experiments

Rats were anesthetized with isoflurane (2% at 1.2 L / minute) and placed in a stereotaxic frame (LPC, France) 30 minutes following saline or PPX injection. Analgesia was performed with ketoprofen (10 mg/kg/day, i.p.) and locally with 2% xylocaine gel.

#### Orbitofrontal activity recording

Juxtacellular in vivo recordings were performed with a glass microelectrode (PG150-T, Harvard Apparatus, Holliston, MA, USA) filled with 0.4 M NaCl solution containing 2% Chicago Sky Blue dye (Sigma-Aldrich, CAS 2610-05-01). Signal was amplified (AxoClamp 900A, Molecular Devices), filtered (low pass: 300Hz; high pass: 10kHz) and digitized (Micro 1401 mk II, Cambridge Electronics Design, England). The recording electrode was lowered into the OFC to screen the spontaneous activity. For each rat, from 5 to 9 descents of 3 mm were done, starting at the coordinates: AP: +3.4 ; ML: +1.8: DV: -4 to -7 (in mm from bregma and dura). Following descents were then carried out to go around the starting point, by moving the electrode 0.2 mm on the mediolateral or anterio-posterior axis. Each neuron was recorded for at least 120 seconds, using Spike2 software (Cambridge Electronic Design Limited, Milton, Cambridge, United Kingdom).

The overall activity of the neuronal population was assessed by measuring the number of spontaneously firing neurons per mm for each rat. Spike and firing pattern were analysed offline, using Spike2 and NeuroExplorer softwares. Spike duration was measured between the beginning of the depolarisation and the end of hyperpolarization. Spike firing patterns were analyzed using NeuroExplorer burst analysis (maximum interval to start a burst = 0.17 seconds, maximum interval to end a burst = 0.3 seconds, minimum interval between bursts = 0.2 seconds, minimum duration of a burst = 0.01 seconds and minimum number of spikes in a burst = 3) as previously described [20]. Neurons that did not present any burst (0% spikes in burst) were considered as non-burtsy neurons, other neurons were considered as bursty neurons.

#### Frontostriatal plasticity recording

A bipolar concentric stimulating electrode (SNEX-100, Rhodes Medical Instruments, USA) was placed in the OFC (AP: +3; ML: +2; DV: -5, in mm from bregma and dura) and a glass microelectrode (PG150-T, Harvard Apparatus, Holliston, MA, USA) filled with 0.4 M NaCl solution was lowered in the DMS (AP: +0.5; ML: +2.75; DV: -4.5 to -6, in mm from bregma and dura). OFC simulations (1 pulse, 600µs, 1mA) were applied every 3 second to evoke a spike in striatal neurons. DMS neurons response to OFC stimulation was recorded using Signal 6 software (Cambridge Electronic Design Limited, Milton, Cambridge, United Kingdom) during baseline and after High Frequency Stimulation (HFS) of cortico-striatal pathway. Spike probability was measured as the number of spikes evoked in response to OFC stimulation over 5 minutes (100 repetitions) and expressed in percent. OFC stimulation intensity was adjusted to induce 50% spike probability during baseline. HFS (2 trains of 100 pulses at 50Hz, inter-train interval: 10s) was performed as previously described [16] using Spike2 software (Cambridge Electronic Design Limited, Milton, Cambridge, United Kingdom).

Electrophysiological recordings were analysed offline, using Spike2 and Signal6 softwares. As for OFC recordings, spike duration was measured between the beginning of the depolarisation and the end of hyperpolarization.

### Tissue processing and histopathological analysis

#### Brain collection

After electrophysiological recording, electrode placement was verified via electrophoretic ejection of Chicago Sky Blue dye (Sigma Aldrich) at the recording site. Sodium pentobarbital (120 mg/kg, i.p.) was injected prior to an intracardiac perfusion with 200 mL NaCl (0.9%) followed by 200 mL ice-cold paraformaldehyde (4%). Brains were postfixed overnight in paraformaldehyde (4%), cryoprotected in sucrose (20% in H2O) at 4°C and frozen in isopentane at -40°C before storage at -80°C. Serial 50 µm coronal free-floating sections were collected and stored at -20°C in a cryoprotectant solution.

#### Tyrosine Hydroxylase immunochemistry

Sections were washed 3 times in Tris Buffer Saline 1X (TBS, 15 minutes each), sections were incubated 10 minutes in H2O2 solution (S202386-2, Agilent Technologies, Santa Clara, CA, USA) to quench endogenous peroxidases. After 3 washes, an incubation of 90 minutes in a blocking solution containing 3% BSA (bovin serum albumin) and 0.3% Triton in TBS 1X was performed. Sections were then incubated 18 hours at 4°C with mouse anti-TH (1/5000, MAB318, Sigma-Aldrich, Saint Louis, MO, USA) in blocking solution. Sections were washed again 3 times in TBS 1X, and then incubated 1 hour at room temperature (RT) with EnVision HRP system anti-mouse (K400111-2, Agilent Technologies, Santa Clara, CA, USA) in blocking solution. Then, sections were washed again three times in TBS 1X and immunoreactions were revealed with DAB peroxidase substrate (K346811-2, Agilent Technologies, Santa Clara, CA, USA). Finally, sections were mounted on gelatin-coated slides and coverslipped with DePeX. TH positive neurons were counted using the optical fractionator method on every 6^th^ section of the SNc as previously described.[21] Systematic random sampling was performed with the Mercator Pro V6.5 (Explora Nova, La Rochelle, France) software coupled with a Leica 5500B microscope. Following delineation of the SNc with x5 objective, counting was done with x40 objective.

#### Recording site vizualisation

Frontal sections were stained with neutral red and Cresyl violet to highlight the recording site. Slices were incubated 4 minutes in a solution of Triton 0.3% before 2 successive water washes. Slices were placed in a bath of alcohol 50° before an incubation in the dye solution and successive incubations with alcohols. Finally, slices were coverslipped with DePeX (Sigma-Aldrich, Saint Louis, MO, USA) mounting medium.

### RNAscope

The multiplex fluorescence RNAscope in situ hybridation was performed on 50 µm thick paraformaldehyde-fixed rat coronal brain sections (Bregma +1.8mm). RNAscope Multiplex Fluorescent Reagent Kit v.2 (323100, Advanced Cell Diagnostics, Newark, CA, United States) was used according to the manufacturer’s instructions. Briefly, following three consecutive washes in TBS, free-floating sections were treated with hydrogen peroxide for 10 minutes at room temperature and mounted on Superfrost Plus and were dried completely. A hydrophobic barrier was created around section. After hydrophobic boundaries had dried overnight, protease digestion by the protease plus (ACDBio) was carried out during 30 minutes. Following tissue pretreatment, samples were hybridized with specific probes to rat eurotrophic tyrosine kinase receptor type 2 (Probe-Rn-Ntrk2, C2 channel, 317531, ACDBio) and dopamine receptor D3 (Probe-RN-Drd3, C3 channel, 449961, ACDBio) mRNA at 40°C for 2 hours in a HybEZ II oven (ACDBio). A three-step amplification process was performed at 40°C for 30, 30 and 15 minutes respectively followed by an HRP development of channel specific signal at 40°C for 15minutes. Final labeling was realized using 570 Opal fluorophore (FP1488001KT, Akoya Biosciences, Marlborough, MA, USA) for *Ntrk2* mRNA and 690 Opal fluorophore, (P1497001KT, Akoya Biosciences) for *Drd3* mRNA. Nuclei were counterstained with DAPI and sections were mounted using Mowiol (Sigma-Aldrich, Vienna, Austria). Sections were washed 3 × 5 min in wash buffer (ACDBio) at RT between each step of the protocol. In each experiment, a slide was used for negative control incubated with a universal negative control probe targeting the *dapB gene* (320871, ACDBio) and for positive control using the 3-plex positive Rn control probe (320891, ACDBio). For quantification of the intensity of fluorescence, images were acquired with a Zeiss Axio Imager.M2 Apotome microscope (Zeiss, Oberkochen, Germany) at ×10 magnification with ZEN imaging software (Zeiss). To allow for quantitative and qualitative comparisons, standardized settings remained constant.

#### Western blot

Rats received an injection of sodium pentobarbital (120mg/kg, i.p.) prior to an intracardiac perfusion with 200 mL 0.9% NaCl. Brains were retrieved, frozen in isopentane at -45°C and stored at -80°C until use. Regions of interest (dorsomedial striatum and orbitofrontal cortex) were collected using a tissue puncher on 150 µm cryostat sections. Proteins were extracted using 1% sodium dodecyl sulfate (SDS) solution in Tris HCl 0.1M with ethylenediaminetetraacetyl (EDTA) 0.01M and phenylmethylsulfonyl fluoride (PMSF), protease inhibitor, and phosphatase inhibitor cocktails at 1%. Following centrifugation at 13000g for 10 min, the supernatants were collected, and the protein concentrations were determined by Pierce BCA protein assay kit (Thermo Scientific, Rockford, USA). Equal amounts of proteins (30µg) were separated by 7.5% SDS-PAGE and transferred to nitrocellulose membranes (Bio-Rad, Marnes-la-Coquette, France). After blocking membranes in Tris Buffer Solution with Tween-20 0.1 M (TBST) and 5% non-fat milk for 1h30 at room temperature, they were incubated overnight at 4°C with the following diluted primary antibodies: 1/1500 to BDNF, 1/800 to TrkB (Cell Signaling Technology, Leiden, The Netherlands) and 1/8000 to alpha-tubulin (Sigma-Aldrich, Saint Quentin Fallavier, France). After washing membranes in TBST buffer, they were incubated with HRP-conjugated secondary antibodies for 1h at room temperature. Following 3 washes in TBST, Chemiluminescence signal was produced with Immobilon solution (Merck Milipore, Bedford, MA) and measured visualized on a PXi image system. Bands were quantified by densitometry using GeneTools software (Syngene, Cambridge, UK). Protein levels were determined after normalizing with alpha-tubulin.

### Statistical analysis

Data are expressed as mean ± Standard Error of the Mean and analyzed using GraphPad Prism 10 software. Normality was tested with the Kolmogorov-Smirnov test. Data having a Gaussian distribution were analysed using two-way and three-way analysis of variance (ANOVAs) followed by Tukey’s post-hoc multiple comparisons test.

## RESULTS

### Effect of alpha-synuclein and PPX treatment on dopaminergic neurons and D3 receptor expression

We used TH immunohistochemistry staining coupled to stereological quantification to assess the level of dopaminergic loss in each animal group. Viral-mediated expression of alpha-synuclein induced a loss of TH-positive neurons in the SNc (F_1, 72_ = 296.1, p<0.0001) and stereological counts showed a significant loss of dopaminergic cells both in saline (p<0.0001) and PPX (p<0.0001) treated alpha-SYN rats compared to saline or PPX-treated GFP rats (**Fig 1A**).

**Figure 1.**
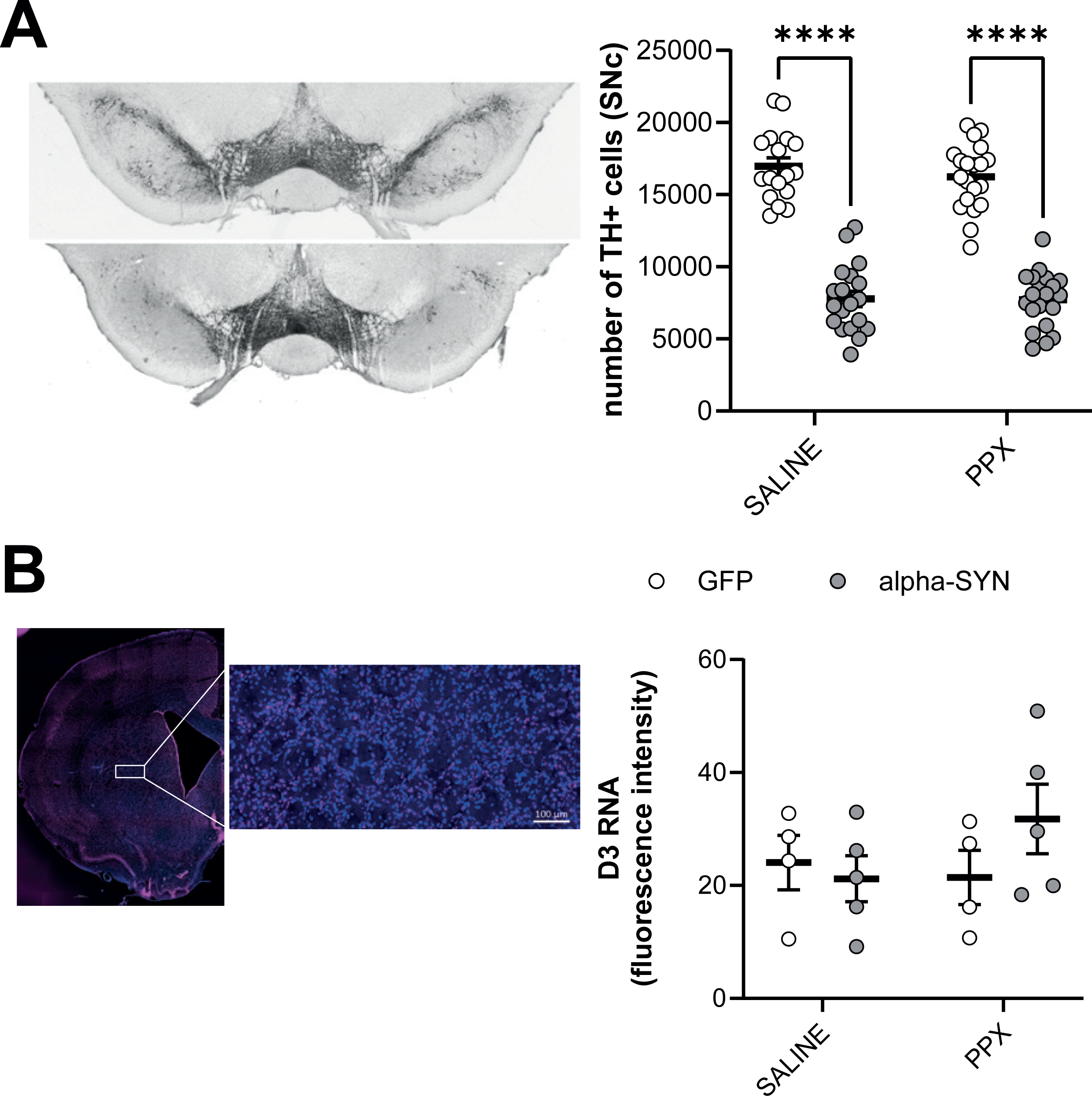
(A) Stereological counts of tyrosine hydroxylase (TH) positive neurons in the SNc and representative images of the SNc in sham (top) and lesioned rats (bottom). GFP SALINE (N=17) ; GFP PPX (N=20) ; alpha-syn SALINE (N=19) ; alpha-syn PPX (N=20). (B) Fluorescence intensity of D3 mRNA in the DMS. GFP SALINE (N=4) ; GFP PPX (N=4) ; alpha-syn SALINE (N=5) ; alpha-syn PPX (N=5). ****p<0.0001; two-way ANOVA followed by Tukey’s post-hoc.

Since impairment of dopamine homeostasis is known to impact dopamine receptors expression in the striatum, we assessed D3 mRNA expression following surgery and PPX treatment using RNAscope. Fluorescence intensity was measured both in the OFC and in the striatum (**Fig 1B**), no signal was observed in the OFC while D3 mRNA levels were detected in the striatum. PPX chronic treatment nor DA neurons degeneration affected D3 RNA levels in the DMS (lesion: F_1, 14_ = 0.53, p=0.48; PPX: F_1, 14_ = 0.59, p=0.45). These results indicate that dopaminergic denervation was successfully induced in the SNc by local viral-mediated expression of alpha-synuclein and that this dopamine cell loss and PPX treatment had no impact on D3 mRNA expression.

### OFC neuronal activity was impacted by dopaminergic lesion and PPX treatment

We investigated the effects of both dopamine denervation and PPX treatment on OFC function by recording OFC neurons using juxtacellular electrophysiology in isoflurane anesthetized rats (**Fig 2A**). To assess the density of spontaneously active neurons, we performed a 3D screen of the OFC (1mm x 1mm x 1mm). As this strategy does not allow for a systematic identification by neurobiotin injection, we thus defined inclusion criteria for neurons as putative pyramidal neurons based on spike duration (**Fig 2A, B**). Out of 737 recorded neurons, 668 neurons whose spike duration was comprised between 0.5 and 1.4 ms were included in the study as putative pyramidal neurons. In average, one spontaneously firing neuron was recorded per mm in all groups (**Fig 2C**) indicating that the overall neuronal activity was not impaired by alpha-synuclein induced dopaminergic loss nor by chronic PPX treatment (lesion: F_1, 25_ = 0.036, p=0.85 ; PPX: F_1, 25_ = 0.002, p=0.96).

**Figure 2.**
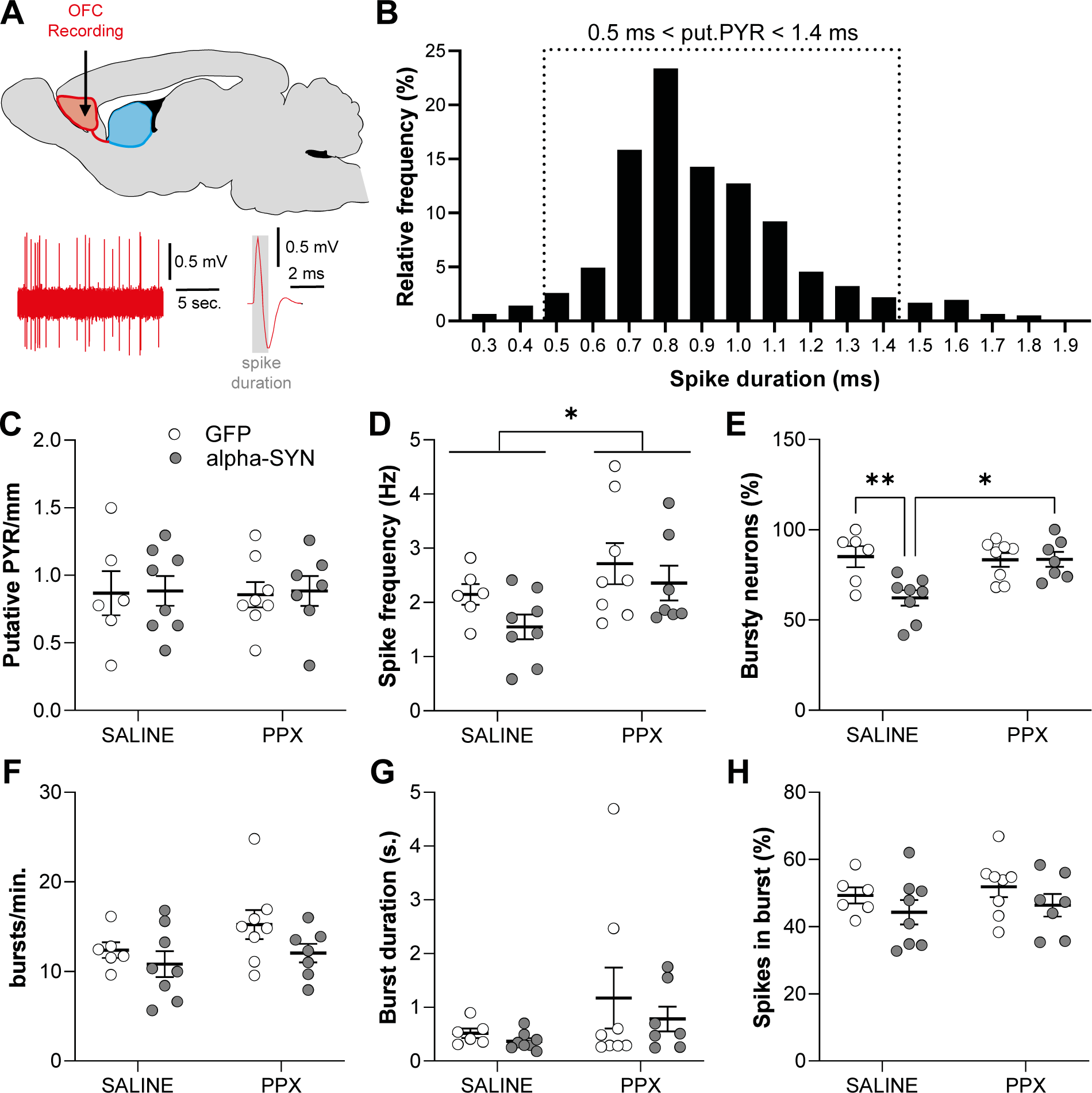
(A) Representative electrophysiological trace of an OFC neuron, the inset represents the action potential shape (averaged over 2 minutes recording), the action potential duration is measured between the two dashed lines. (B) Distribution of spike durations (ms), the dashed line indicates the limits of inclusion criteria for putative pyramidal neurons. (C) Density of spontaneously active putative pyramidal neurons per mm. (D) Spike frequency (Hz). (E) Proportion of neurons exhibiting burst firing activity (%). (F) Bursting frequency (Bursts per minute). (G) Burst duration (seconds). (H) Percentage of spikes in burst. GFP SALINE (n=157 neurons ; N=6 rats) ; GFP PPX (n=162 ; N=8) ; alpha-syn SALINE (n= 191; N=8) ; alpha-syn PPX (n= 158; N=7). *p<0.05 ; **p<0.01 ; two-way ANOVA followed by Tukey’s post-hoc.

We then analysed in more details spike firing patterns by measuring average firing frequency (**Fig 2D**) and burst firing properties (**Fig 2E-H**). Spike firing frequency was increased by PPX treatment (F_1, 25_ = 5.09, p<0.05) but no effect of the lesion nor interaction between lesion and treatment were found, indicating that the dopaminergic denervation had no effect on OFC firing frequency. However, there was a significant effect of alpha-synuclein induced dopaminergic lesion on the proportion of neurons exhibiting burst firing (F_1, 25_ = 6.39, p<0.05), with a decreased proportion of bursty OFC neurons in saline-treated alpha-SYN compared to GFP-rats (p<0.01) and this effect was rescued by chronic PPX treatment (**Fig 2E**). Mean frequency in bursts, burst duration and percentage of spikes in burst were not altered by the lesion nor by PPX treatment (**Fig 2F-H**). These results indicate that alpha-synuclein induced nigral degeneration had no effect on the overall spontaneous activity of OFC neurons, but impacted their firing pattern, as shown by a decreased bursting activity. Interestingly, this effect was rescued by chronic PPX treatment.

### Fronto-striatal plasticity is dramatically impacted by nigrostriatal dopaminergic neurodegeneration and PPX treatment

High frequency stimulation (HFS) was applied in the OFC to induce plastic changes in the connection strength between OFC and DMS neurons (**Fig 3A, B**). Juxtacellular recordings from striatal neurons were performed before and up to 45 minutes after HFS. Three-way ANOVA indicated a main effect of time (F_4.783, 95.66_ = 22.35, p<0.0001), treatment (F_1, 20_ = 23.55, p<0.0001), lesion (F_1, 20_ = 23.38, p<0.0001) and time x treatment x lesion interaction (F_11, 220_ = 8.563, p<0.0001) (**Fig 3C**). As expected [16], HFS induced an increase in spike probability in saline-treated GFP-rats (GFP SALINE pre-HFS vs GFP SALINE post-HFS, p<0.05) while a decrease of DMS response to OFC stimulation was observed following dopaminergic lesion (alpha-SYN SALINE pre-HFS vs alpha-SYN SALINE post-HFS, p<0.0005) and this was not rescued by PPX chronic treatment (alpha-SYN PPX pre-HFS vs alpha-SYN PPX post-HFS, p<0.001). Chronic PPX treatment alone also resulted in a long-term decrease of spike probability following HFS (GFP PPX pre-HFS vs GFP PPX post-HFS, p<0.0001). The OFC-DMS response probability was increased only in control rats, no difference was observed between lesioned, PPX-treated or lesioned and PPX-treated rats. These results indicate that both dopaminergic lesion and chronic PPX treatment induced a reversal of plastic changes measured following HFS compared to control animals and that PPX treatment failed rescuing the effect of dopamine denervation on this parameter.

**Figure 3.**
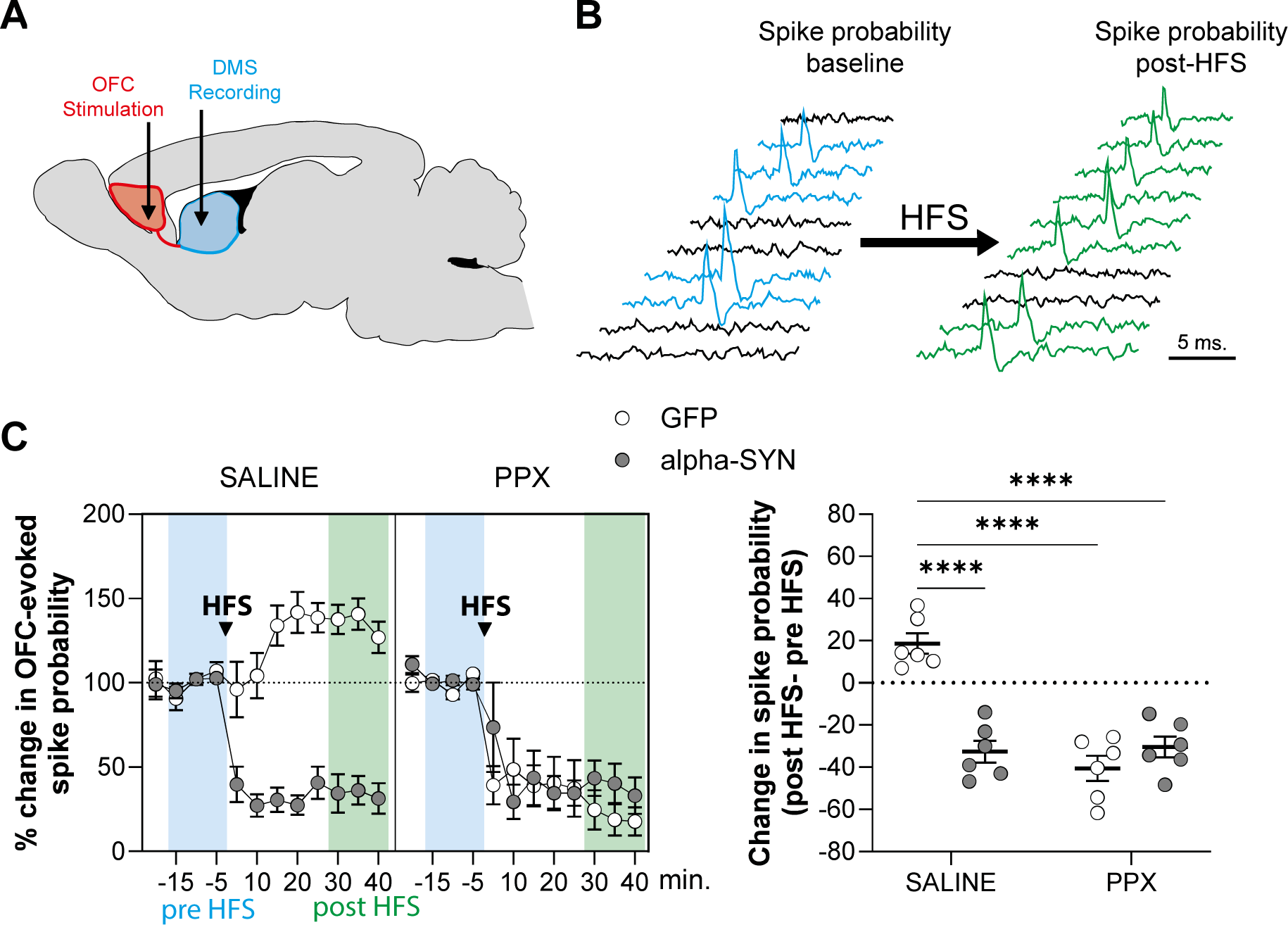
(A) Representative electrophysiological trace of spike responses evoked in a DMS neuron by OFC stimulation. (B) Black traces represent trials that do not exhibit spike responses to OFC stimulations and colored traces represent trials exhibiting spike response to OFC stimulation both before (baseline, blue) and after (post HFS, green) high frequency stimulation of the OFC-DMS pathway. Spike probability was measured as the number of responding trials over 5 minutes. (C) Spike probability (% of baseline) 15 before and 30-45 minutes after HFS. Delta change of response probability after HFS compared to baseline (%). GFP SALINE (N=6) ; GFP PPX (N=6) ; alpha-syn SALINE (N=6) ; alpha-syn PPX (N=6); ****p<0.0001 ; two-way ANOVA followed by Tukey’s post-hoc.

**Figure 4.**
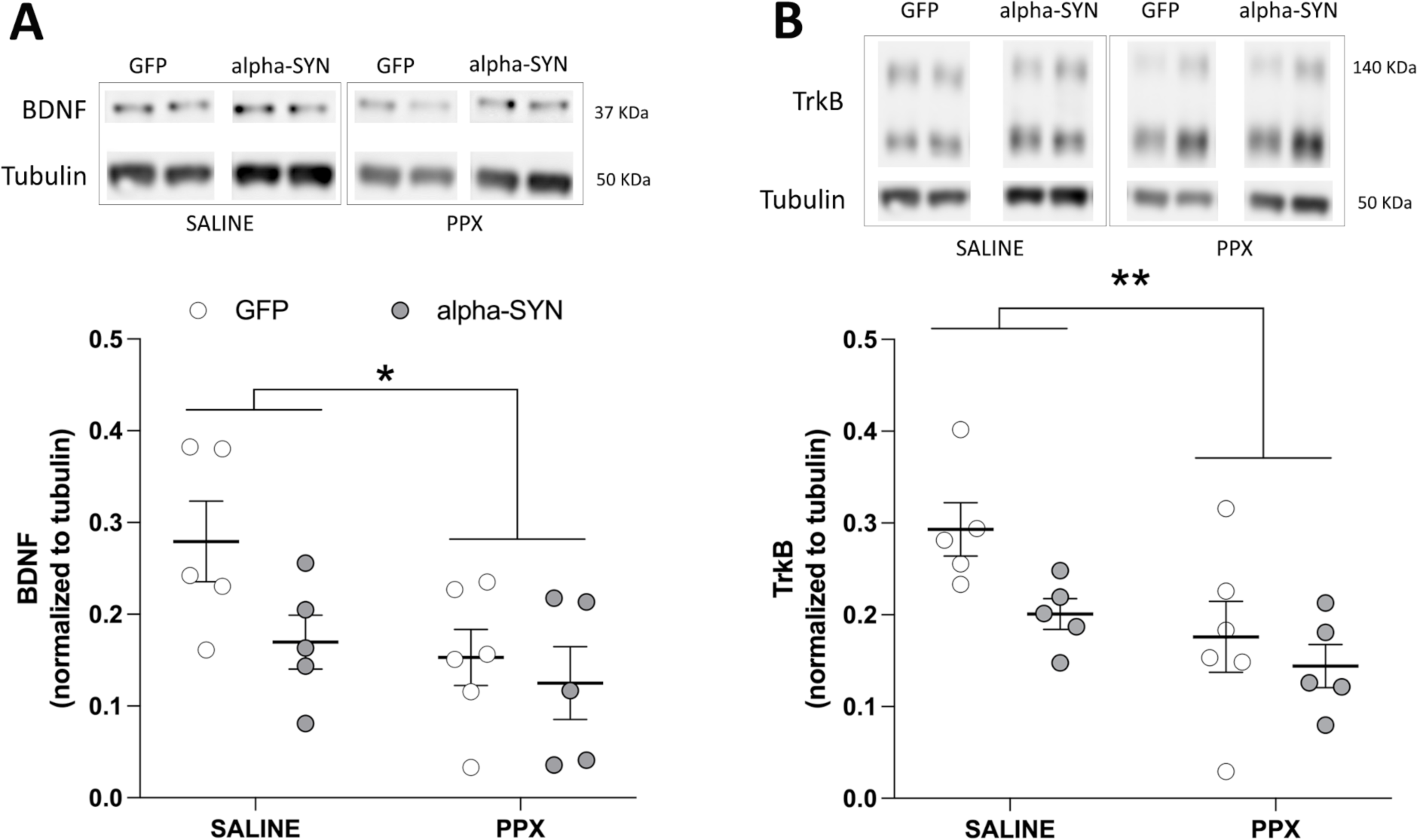
(**A**) BDNF immunoblot levels and (B) TrkB immunoblot levels in the DMS. GFP SALINE (N=5) ; GFP PPX (N=5) ; alpha-syn SALINE (N=5) ; alpha-syn PPX (N=6). *p<0.05 ; **p<0.01 ; two-way ANOVA followed by Tukey’s post-hoc.

### PPX treatment leads to altered striatal BDNF/TrkB levels

To probe potential mechanisms involved in the alteration of OFC-DMS plasticity following alpha-synuclein induced dopaminergic neurodegeneration or PPX treatment, we assessed the levels of BDNF and TrkB in the DMS by western blot. Nigrostriatal lesion induced a trend for a decrease of striatal BDNF levels (F_1, 17_ = 3.64, p=0.07). This decrease was further exacerbated in all groups treated with PPX (F_1, 17_ = 5.62, p<0.05). Similarly, there was a trend for reduced levels of full-length TrkB following nigrostriatal degeneration (F_1, 17_ = 4.35, p=0.052), which were further decreased in all animals exposed to PPX (F_1, 17_ = 8.53, p<0.01). Levels of truncated TrkB were unaffected by the lesion or PPX (data not shown).

## DISCUSSION

Motor basal ganglia loops have been extensively studied following dopamine denervation and dopamine replacement therapy with L-dopa [14]. However, little is known regarding associative circuits and the effects of PPX. We hypothesized that dopamine denervation and chronic dopamine agonist administration could induce OFC-DMS dysfunction paralleling those described in motor loops. To this end, we investigated the respective contributions of nigrostriatal dopaminergic neurodegeneration and chronic treatment with the D2/D3 agonist PPX on the OFC-DMS pathway in a rat model of PD.

Here, we highlight an impairment of associative basal ganglia loops in a viral-mediated model of PD. Indeed, we demonstrated that OFC putative pyramidal neurons exhibited a decrease in burst firing following dopamine denervation in the SNc. However, it has to be noticed that this impairment of OFC activity remains restricted to this particular firing pattern and that the mean firing activity did not seem to be impacted. This effect of DA lesion on OFC neurons seem to differ from that reported in the motor cortex[22], but studies conducted in the motor cortex from rats showed diverging results. In freely moving rats as [8] well as in urethane anesthetized rats [23], 6-hydroxydopamine (6-OHDA)-induced nigrostriatal DA lesion resulted in decreased pyramidal neurons firing rate while in isoflurane anesthetized rats 6-OHDA treatment induced a marked increase in firing rate and bursting pattern of OFC pyramidal neurons [24]. Such discrepancies may be related to differences in the types of DA denervation (medial forebrain bundle vs intranigral 6-OHDA injections) and anaesthetics. Overall, our results indicate that progressive bilateral nigro-striatal DA denervation had no effect on the mean firing rate from OFC neurons but decreased their bursting activity. This latest consequence of DA cell loss was rescued by the D2/D3 agonist, PPX.

While the role of dopamine receptors has been well studied in the medial prefrontal cortex (mPFC) [25], there is little information on how dopamine modulates pyramidal neurons in the OFC. Our results showing enhanced putative pyramidal neuron activity after D2 receptor activation are consistent with the finding that quinpirole acting on D2 receptors increases the excitability of mPFC pyramidal neurons in adult mice [26]. Only a few studies performed *ex vivo* on mice OFC neurons indicate that DA bath application decreases pyramidal neurons excitability [27] through D2 receptor activation [28]. Our results, obtained from *in vivo* recordings, indicate however that in control animals D2/D3 receptor activation by PPX induces an increase in putative pyramidal neurons firing both in lesioned and in non-lesioned rats. The discrepancy between our data and previously reported effect of D2 receptor activation on pyramidal OFC neurons can be due to the differences in species and preparation. Indeed, in anesthetized animals the network is intact resulting in preserved local and distal inputs. Thus D2/D3 receptor effect in our conditions could also be indirect through local GABAergic interneurons inhibition [26, 29].

The modest effect observed in this PD model on OFC function was expected since OFC dopamine innervation mostly comes from ventral tegmental area dopamine neurons [5, 27] that are preserved in this model [17]. Although discrete with regards to the overall neuronal activity, a change in burst firing might account for meaningful alteration of information processing.

The efficacy of OFC to DMS connection was assessed by measuring the efficacy of OFC electrical stimulation to evoke action potential responses in DMS neurons. Striatal projection neurons are medium sized spiny neurons (MSNs) that represent 95% of the striatal neurons and are classified according to their projection target. Striatonigral neurons project directly to the substantia nigra pars reticulata and express D1 dopamine receptors while striatopallidal neurons form the indirect pathway and express D2 receptors [30]. Mallet et al. (2006) have shown in an elegant study the differential modulation of both MSN populations in the dorsolateral striatum following unilateral 6-OHDA lesion [31]. Dopamine cell loss inhibited the direct pathway whereas indirect pathway neurons were activated. Even though we were not able to identify their D1 or D2 phenotype, all recorded DMS neurons were similarly affected by the progressive DA loss and by PPX treatment.

Our results demonstrate that the efficacy of this pathway is increased following high frequency stimulation in control rats as expected [16], while the same stimulating protocol induced a decrease in the OFC-DMS efficiency following dopamine denervation, PPX treatment and the combination of both DA loss and PPX. These results are echoing those reported by the extensive work conducted in the motor cortico-striatal loop to decipher the synaptic mechanisms of L-dopa induced dyskinesias following dopamine cell loss. Synaptic plasticity is differentially altered following loss of DA homeostasis. Partial DA denervation can impact either LTD or LTP while extensive denervation impairs both forms of plasticity (Schirinzi et al., 2016). Accordingly, the progressive and partial nigro-striatal DA lesion in our study resulted in a loss of LTP, and a switch to LTD. Moreover, while L-dopa treatment restores the loss of LTP in the motor cortico-striatal pathway [32], our data show no such recovery following PPX treatment in the associative loop. Eventually, PPX itself induced a switch from LTP to LTD in control rats, similar to that observed after DA lesion. This effect the D2/D3 agonist is coherent with the promotion of LTP by D2 antagonist haloperidol [33].

In addition to its potent effects on neuronal survival, function and dendritic spine morphology, BDNF plays a key role in synaptic plasticity and LTP [34, 35]. To probe for potential mechanisms associated with the alterations of frontostriatal plasticity observed following alpha-synuclein induced nigral neurodegeneration and PPX treatment, we assessed BDNF and TrkB levels in the DMS. In PD, BDNF levels are reduced in the substantia nigra, caudate and putamen [36]. In lesioned rats, there was only a trend for decreased BDNF levels in the DMS. Consistent with a predominant cortical, and to a lesser extent nigral origin of striatal BDNF [37], this marginal effect on BDNF levels may be attributable to the fact that the loss of nigral dopaminergic neurons occurring in our model is only partial. Interestingly, chronic treatment with PPX significantly decreased DMS BDNF levels in both sham and lesioned rats with no additive effect of the lesion, thus indicating a main contribution of PPX itself. These results extend previous findings showing that a high dose of PPX (1 mg/kg twice a day) can reduce BDNF mRNA levels in the habenula in normal rats and in the amygdala and nucleus accumbens in 6-OHDA lesioned rats [38].

BDNF binding with high affinity to TrkB results in downstream activation of the mitogen-activated protein kinases and extracellular signal-regulated kinases signaling pathways that are key effectors of post-synaptic LTP [35]. Similar to BDNF, nigral neurodegeneration led to a trend for decreased TrkB levels in the DMS. PPX treatment had a more pronounced effect, leading to significantly reduced levels of TrkB in both sham and lesioned rats. Similar effects of PPX on TrkB mRNA levels were found in the striatum of control and 6-OHDA-lesioned rats [38]. Given the demonstrated role for BDNF and TrkB signaling in mediating LTP [34, 35], our results suggest that alterations in BDNF and TrkB may contribute to the alterations of frontostriatal plasticity observed in our study.

## CONCLUSION

Our results suggest that nigral dopaminergic loss and chronic PPX lead to both pre and post synaptic alterations in the OFC-DMS pathway. Altered OFC activity may affect the encoding of stimulus reinforcement association learning [39] and impaired frontostriatal plasticity may impact the changes in synaptic efficacy that are necessary for optimal responses and behavioural adaptations. Such frontostriatal dysfunction precluding adequate information processing in associative basal ganglia loops may contribute to the expression of NMS and create a vulnerability state for the emergence of cognitive symptoms following dopamine replacement therapy in suceptible individuals.

## ACKNOWLEDGMENTS

The Université de Poitiers and the Institut National de la Santé et de la Recherche Médicale provided infrastructural support. This work was supported by France Parkinson (grant #R20006GG/RAK20003GGA to MBM and post-doctoral fellowship to MD), région Nouvelle Aquitaine CPER 2015-2020 and FEDER 2014-2020 programs. The authors thank Eric Balado and Catherine Le Goff for technical support. This study has benefited from the facilities and expertise of PREBIOS core facility at the University of Poitiers. The funders had no role in study design, data collection and analysis, decision to publish, or preparation of the manuscript.

